# Long telomere inheritance through budding yeast sexual cycles

**DOI:** 10.1101/2025.06.05.658123

**Authors:** Vasilisa Sidarava, Sarah Mearns, David Lydall

## Abstract

The ends of linear eukaryotic chromosomes are protected from being recognized as DNA double-strand breaks by telomeres, containing repetitive DNA sequences which bind specific proteins. In humans, mutations in telomere regulatory genes lead to short or long telomere syndromes. These syndromes often show genetic anticipation, where disease has earlier onset and a more severe manifestation in each new generation. Later generations inherit not only the mutation affecting telomere length, but also abnormal length telomeres.

Many aspects of telomere length homeostasis are conserved between mammals and yeast. Here we explored telomere length inheritance patterns through the sexual cycle in yeast. Analysis of single telomeres, rather than bulk telomeres, shows that if haploid yeast with short telomeres mate with wild-type yeast creating diploids, short telomere lengths rapidly normalize (within 30 cell divisions). However, long telomeres inherited from one parent can persist for more than 200 mitotic cell divisions. Long telomere can also be transmitted through more than one round of meiosis, independently of mutations that cause long telomeres. These patterns, along with haploinsufficiency effects, show that even in yeast there is a complex relationship between telomere length, telomere length inheritance, and mutations that affect telomere length. Our findings may have implications for families affected by telomere syndromes.

**Article summary:** Telomeres protect the ends of chromosomes from DNA damage responses. In humans, mutations causing short or long telomeres, lead to inherited disease syndromes, often showing evidence of genetic anticipation. Here we show that in budding yeast, long telomeres can be transmitted through several generations in the absence of mutations causing the long telomere phenotype. Therefore, even in a simple single-celled eukaryotic organism, complex patterns of telomere length inheritance occur.

**Graphical abstract:** 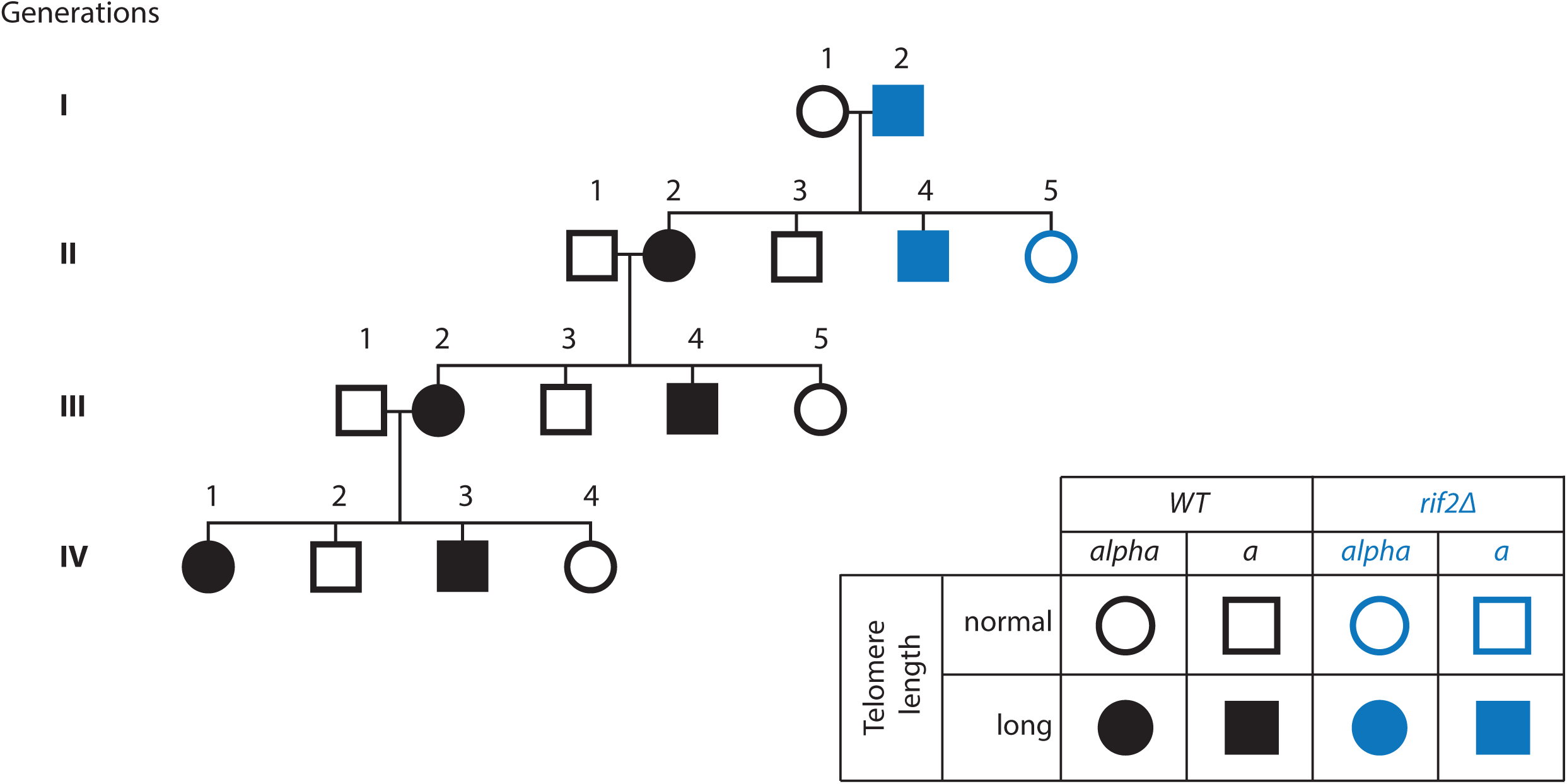

## Introduction

Telomeres ensure that linear chromosome ends do not behave like double strand break (DSB) ends (Blackburn 1991; Pfeiffer and Lingner 2013). Whereas internal DSBs are potent inducers of the DNA damage response (DDR), telomeres are not. In most eukaryotes, telomeres comprise an array of short repetitive DNA sequences and associated proteins and RNA. The DNA repeats (TG_1-3_ in budding yeast, TTAGGG in humans) are largely double-stranded, ending with a short single-stranded 3’-overhang (G-tail) (Greider 1996) that can form a t-loop (Doksani et al. 2013). Telomeric DNA is bound by multiple “capping proteins”, which help prevent chromosome ends from stimulating the DDR (Takai et al. 2003).

The end replication problem, that DNA polymerases cannot fully replicate the ends of linear DNA molecules, applies at chromosome ends. While most of the telomeric DNA is replicated in a semi-conservative manner during S phase, the very end of the 3’ overhang cannot be fully copied by DNA polymerases. To compensate for this end replication problem, most eukaryotic cells, including yeast and human cells, employ telomerase, to add telomeric repeats at the 3’ ssDNA telomeric termini using its intrinsic RNA template (Greider and Blackburn 1985; Yu et al. 1990). Since the addition of terminal telomeric DNA to chromosome ends is not a semi-conservative process, telomerase activity needs to be tightly regulated. If telomerase were insufficiently active, chromosome ends would shorten and stimulate a DDR (Enomoto et al. 2002; IJpma and Greider 2003). The existence of mechanisms to rapidly shorten overly long telomeres (telomere rapid deletion) (Li and Lustig 1996; Pickett et al. 2009) suggests that long telomeres are a problem as well. Natural selection presumably determines optimal telomere length for each species.

Control of telomere length is complex. In budding yeast, more than 300 different genes have been shown to affect telomere length (Askree et al. 2004). In humans it is likely that telomere length regulation is at least as complicated. In somatic tissues, telomerase gene expression is developmentally downregulated (Wright et al. 1996; Ulaner et al. 1998), which is thought to limit cell proliferation and to protect against cancer (Shay 2016). In addition, in mammals, different cell types can replicate and maintain telomeres in different ways. For example, while the capping protein TRF2 is essential for telomere protection in somatic cells, stem cells depend less on this protein (Markiewicz-Potoczny et al. 2021).

It is well established that a number of heritable syndromes are associated with incorrect telomere lengths (Armanios 2013). Broadly, in short telomere syndromes, it is thought that the short telomeres stimulate DDRs that inhibit cell division and cause senescence and/or apoptosis, compromising tissue function. This can cause pulmonary fibrosis, liver disease or bone marrow failure (Armanios 2013). In long telomere syndromes, which are less well studied, long telomeres appear to facilitate cellular transformation by permitting cell proliferation, and are associated with melanoma, chronic lymphocytic leukemia, and papillary thyroid cancer (McNally et al., 2019; DeBoy et al. 2024; Savage 2024).

Understanding how different genes affect disease in patients with telomere syndromes is particularly challenging partly because of the phenomenon of genetic anticipation. Genetic anticipation occurs because abnormal telomeres, as well as mutations causing abnormal telomeres, are transmitted through the generations (Armanios et al. 2005). For example, the effect of a mutation in a grandparent, with normal telomeres, may be less than in a grandchild, who has inherited the identical mutation but done so in combination with short telomeres.

In humans the telomeres inherited by egg or sperm cells are the products of a large number of mitotic and meiotic cell divisions during development and differentiation. It seems likely that telomere length in oocytes or sperm cells is largely determined by number of cell divisions the cell progenitors have undertaken. Human females generate oocytes *in utero* and it is estimated that around 30 cell divisions are needed to produce oocytes (zygote to oocyte) (Drost and Lee 1995) (Figure 1A). Males generate sperm throughout adult life, with the number of cell divisions any sperm cell has undergone depending on the male’s age. It is estimated that sperm cells from thirty year-old males will have undergone around 400 cell divisions (zygote to sperm cell) (Drost and Lee 1995) (Figure 1A). Although human oocyte and sperm cells undergo very different numbers of cell divisions before being generated, it seems that telomeres in sperm of older men are longer than those in younger men (Allsopp et al. 1992; Hjelmborg et al. 2015; Eisenberg and Kuzawa 2018), suggesting it is difficult to directly correlate the number of cell divisions with telomere length.

**Figure 1.**
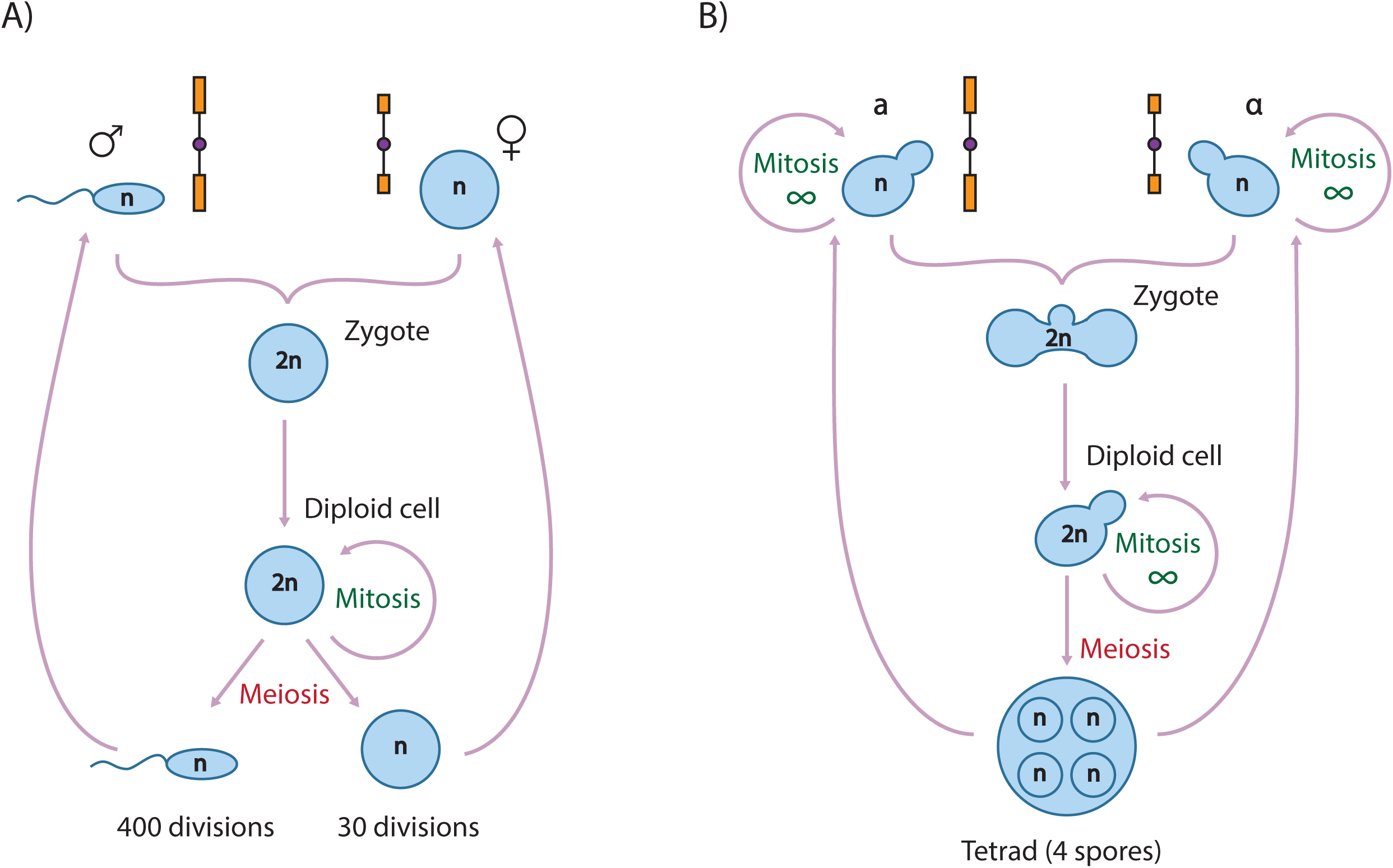
Human and budding yeast sexual cycles. A) In males, sperm cells are produced throughout life, and for 30-year-olds, the number of divisions between a zygote and a sperm cell is estimated to be 400. Females are born with their egg cells, which had been produced *in utero*. The number of divisions between a zygote and an egg cell is around 30. B) In yeast, the number of mitotic divisions in haploids and diploids is unlimited. Zygotes are illustrated inheriting chromosomes with wild-type and short telomeres (telomeres, blue rectangles; centromeres, purple circles).

In yeast, it is well established that telomere length is regulated during the cell cycle. Telomerase is active towards the end of S phase and telomere rapid deletions also occur during this phase of the cell cycle (Taggart et al. 2002). Since the yeast sexual cycle can be used to model aspects of the human sexual cycle (Figure 1B) we wondered if yeast might be used to model the extent to which eukaryotic telomere length changes are influenced by the number of mitotic and meiotic cell divisions. Our experiments show that even in yeast there are complex patterns of telomere length inheritance through the sexual cycle.

## Materials and methods

### Yeast strains

Strains used are listed in Table 1. *RIF1* and *RIF2* deletions in the W303 background were made by transforming DLY3001 and DLY8460 with a PCR-amplified G418 resistance cassette from *rif1Δ* and *rif2Δ* strains of the S288C background, respectively.

**Table 1.**
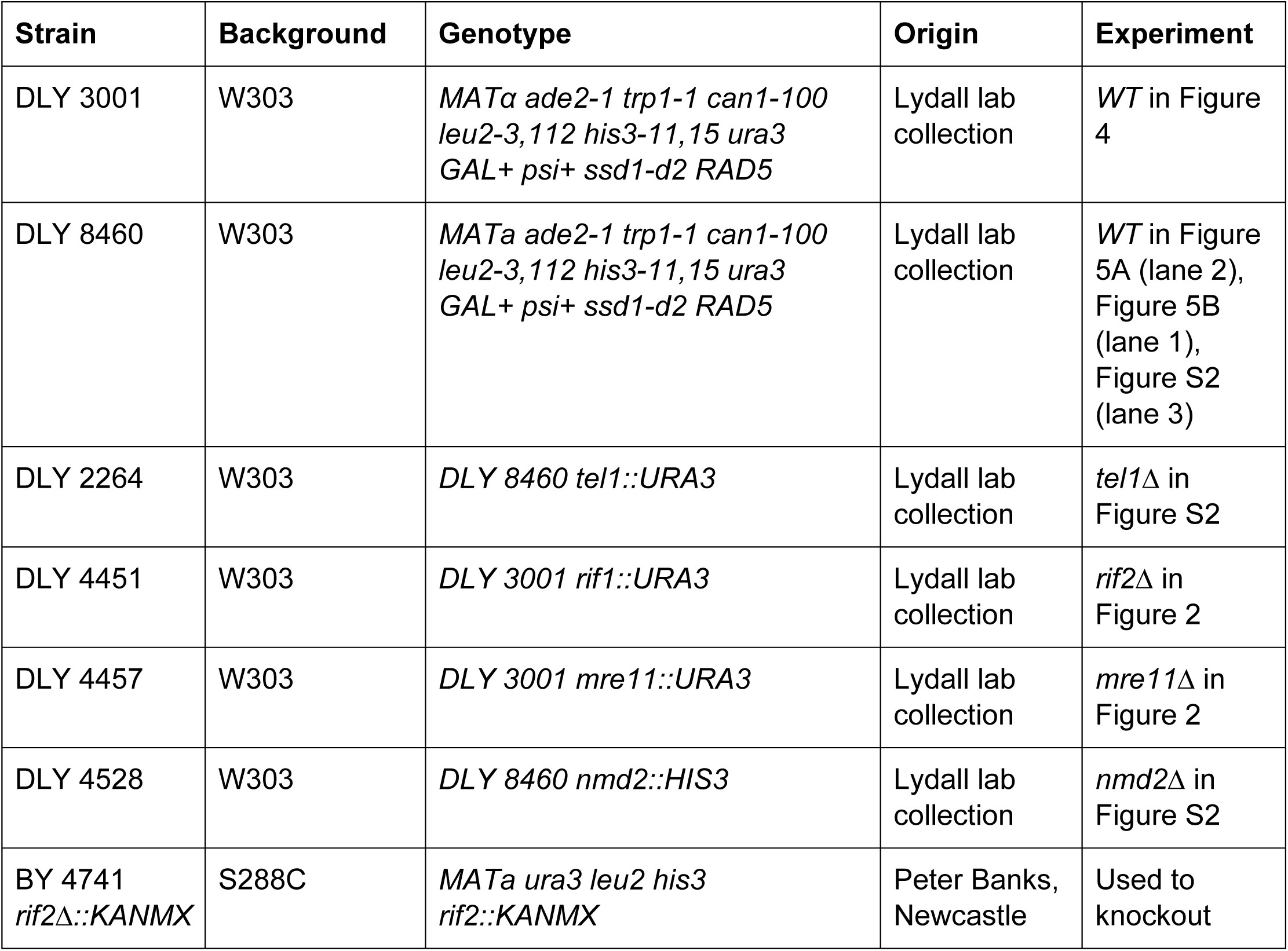
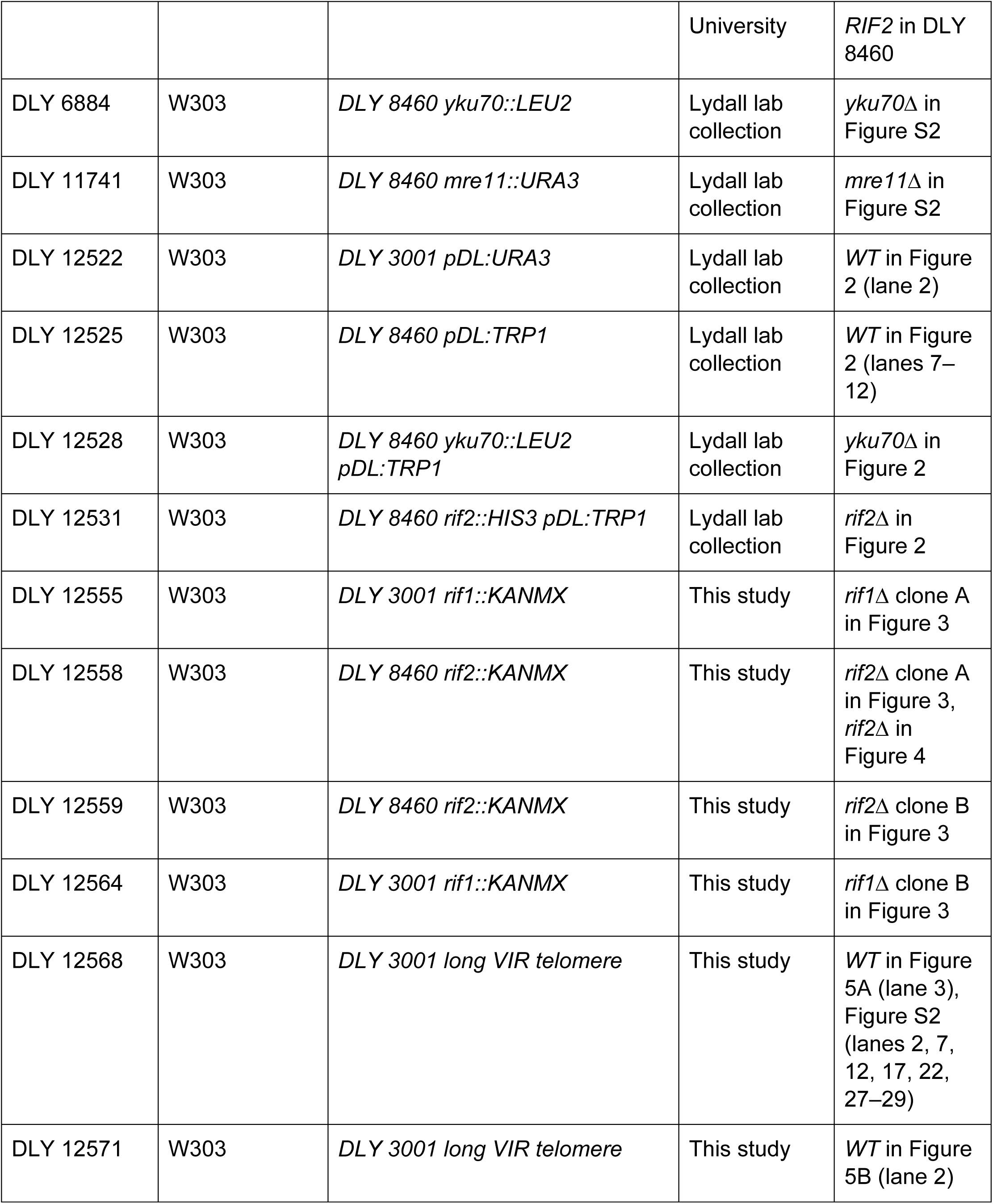
Strains used in this study.

### Yeast propagation and genetic manipulation

Yeast were routinely cultured at 30°C on YEPD plates (1% yeast extract, 2% peptone, 2% agar, 2% dextrose) supplemented with adenine (75 mg/l).

Haploid transformants were selected on YEPD-G418 plates, then streaked for single colonies on YEPD-G418 plates, and patched on YEPD plates. Haploid clones were passaged every 3.5 days by streaking for single colonies. To produce diploids, haploid strains were mixed on YEPD plate and mated. When auxotrophic selection was possible, diploids were selected by plating mating mixtures on doubly selective media. Alternatively, zygotes were picked after mating mixtures were incubated for 3 hours at 30°C. Zygotes were isolated using a tetrad dissection microscope. Colonies derived from zygotes were cultured for 3.5 days before being passaged. Diploid yeast clones were passaged every 3.5 days by streaking for single colonies. For sporulation, diploids were grown to saturation in liquid YEPD on a wheel at 30°C and incubated for 3 days on a wheel at 23°C in liquid sporulation medium (1% KAc and the following supplements (per L): 1 g yeast extract, 0.5 g dextrose, 0.1 g supplement powder mix (2g histidine, 10 g leucine, 2 g lysine, 2 g uracil, 2 g tryptophan, 3 g adenine)). Tetrads were dissected on YEPD plates, and spore progeny were grown for 3.5 days. Prior to DNA extraction and telomere length analysis, the cells went through different numbers of cell divisions, depending on the cell’s origin (Table 2).

**Table 2.** Cell division numbers.

| Cell origin | In a colony | In a patch | In liquid culture | Reference |
| --- | --- | --- | --- | --- |
| Auxotrophically selected diploids | 25 (from 1 to $2.8 \times 10^7$ cells) | 4 (from $1.4 \times 10^7$ to $1.6 \times 10^8$ cells) | 3 (from $5 \times 10^7$ to $2.9 \times 10^8$ cells) | P1 diploids in Figures 2, S1 |
| All diploids from P2 and beyond | 25 (as above) | 4 (as above) | 3 (as above) | Diploids in Figure 2, S1, and S2 (except P1); |
| Haploid transformants | 26 (1<sup>st</sup> G418 plate: from 1 to $4.6 \times 10^7$ cells) and 23 (2<sup>nd</sup> G418 plate: from 1 to $6.8 \times 10^6$ cells) | 5 (from $6.8 \times 10^6$ to $1.6 \times 10^8$ cells) | 3 (est. from data for haploids) | P1 haploids in Figure 3 |
| Haploids from P2 and beyond | 26 (from 1 to $7.9 \times 10^7$ cells) | 3 (from $2.0 \times 10^7$ to $15.3 \times 10^7$ cells) | 3 (from $5.1 \times 10^7$ to $3.6 \times 10^8$ cells) | Haploids in Figure 3 (except P1) |
| Pulled zygote | 25 (from 1 to $2.8 \times 10^7$ cells) | | 4 (from $1.4 \times 10^7$ to $1.8 \times 10^8$ cells) | Diploids in Figures 4, 5; P1 in Figure S2 |
| Germinated spore | 25 (from 1 to $2.4 \times 10^7$ cells) | | 4 (from $1.2 \times 10^7$ to $2.1 \times 10^8$ cells) | Haploid progeny in Figures 4, 5 |

**Table 3.**
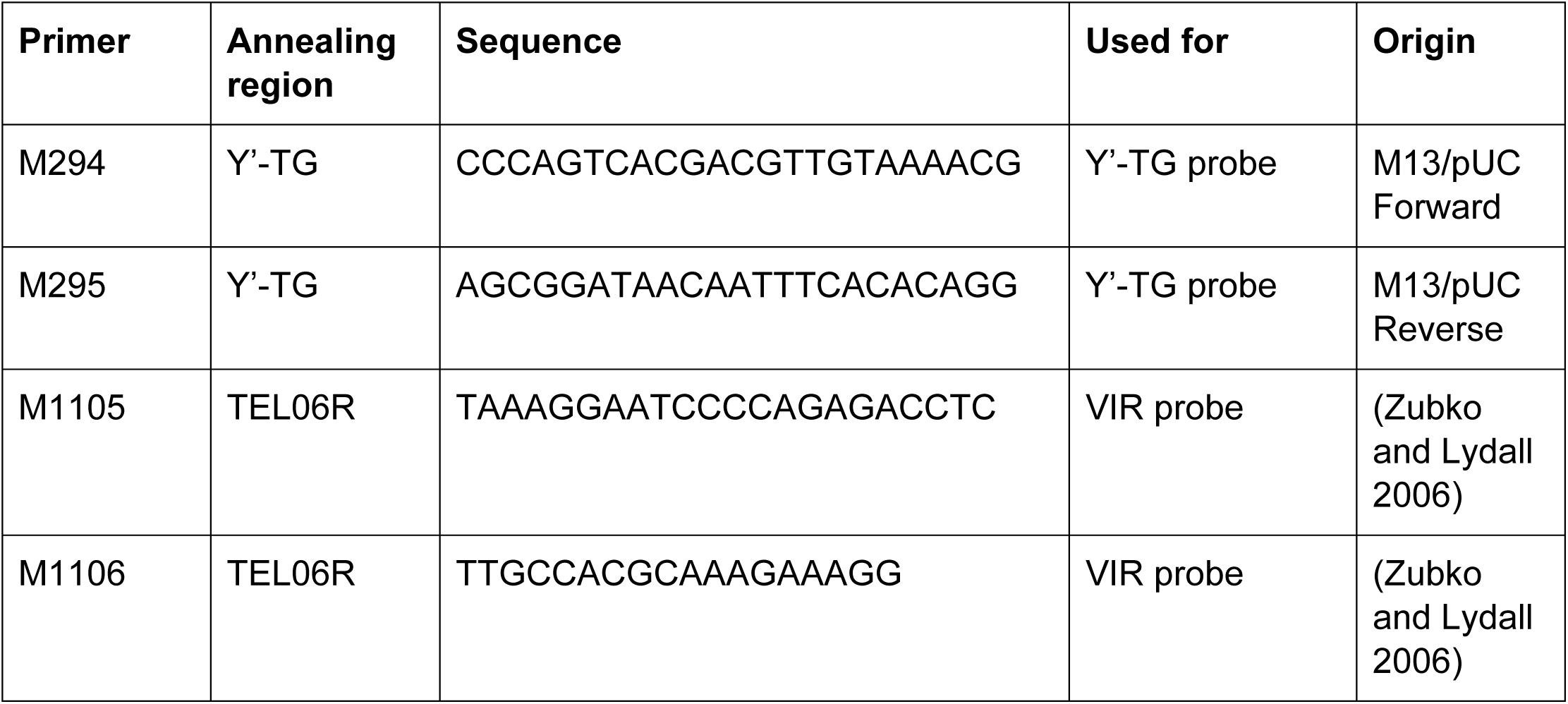
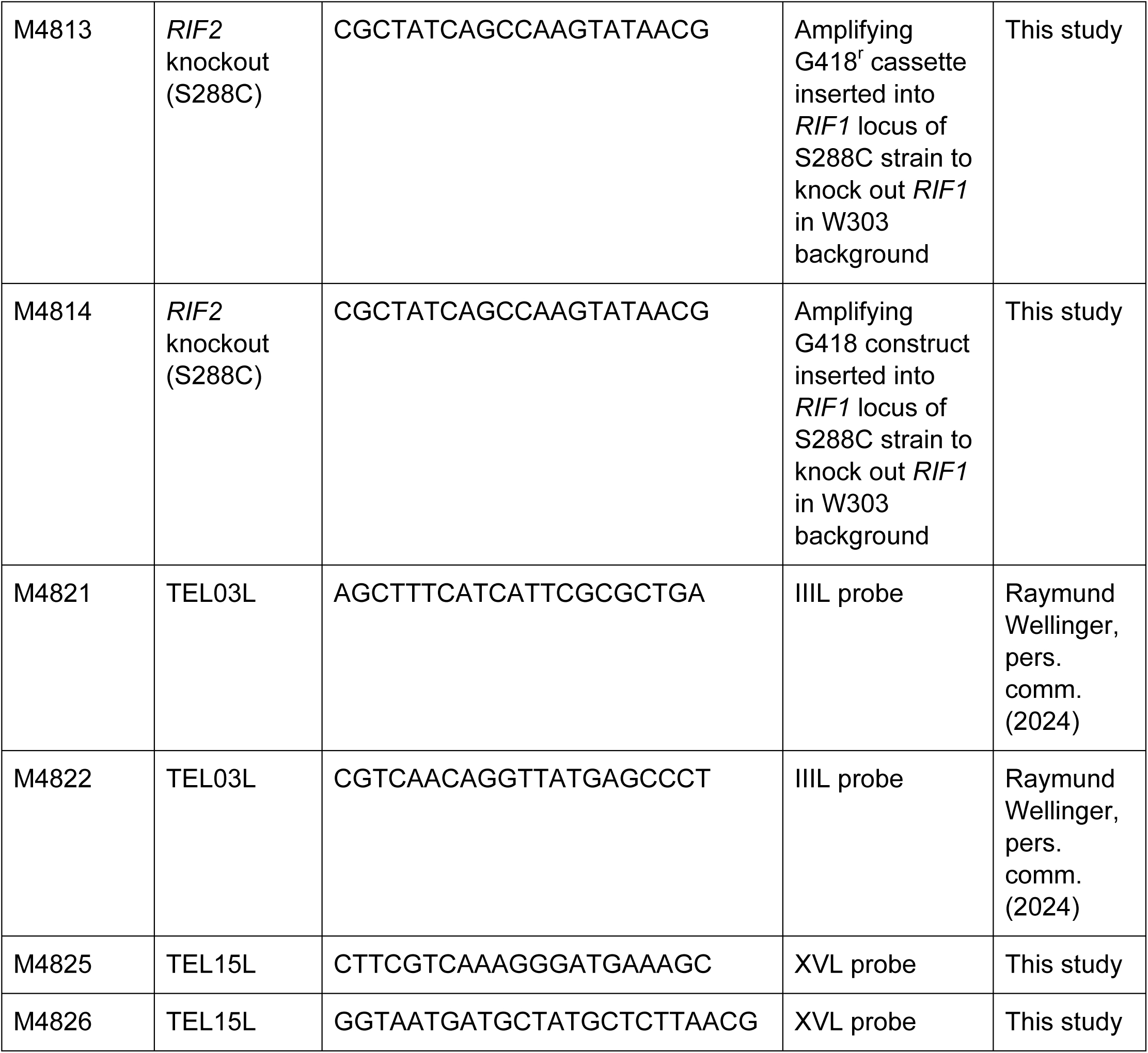
Primers used in this study.

### Estimation of yeast cell divisions

The number of cells divisions taking place during culture (colony, patch, or overnight liquid) was estimated using the formula: 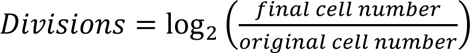. We assumed a colony began as single cell (original cell number = 1). The final cell number for each culture was determined by resuspension or dilution in water and counting in a hemocytometer. In all cases four independent colonies/cultures were used to estimate mean cell numbers. Half a colony was typically used to make an overnight patch, and one-third of the patch was used to inoculate a 2 ml liquid culture. If the patching step was skipped, half a colony was used to inoculate a 2 ml liquid culture.

### Telomere length detection

Genomic DNA was extracted and digested with *Xho*I (Y’-TG and VIR blots) or *Ase*I (XVL and IIIL blots) for 5 hours at 37°C. DNA samples were run on a 1% agarose gel at 17 volts for 16 hours. Gel transfer, blotting, probe hybridization and detection were performed according to manufacturer’s instructions using DIG High Prime Labelling and Detection Starter Kit II (Roche). Probes were synthesized using PCR DIG Labeling Mix (Roche). VIR, XVL and IIIL probes were amplified from DLY 12525 genomic DNA using primers M1105, M1106; M4825, M4826; and M4821, M4822, respectively. The Y’-TG probe was amplified from pDL678 (pHT128, pYtel) (Tsubouchi and Ogawa 2000) using primers M294 and M295. Telomere length was estimated using ImageLab software (version 6.0.0, Bio-Rad Laboratories).

## Results

### Long telomeres persist in diploids, short telomeres quickly normalize

We first addressed what happens to telomere length in diploid cells that have inherited telomeres of differing lengths from their haploid parents. Haploid yeast with normal (N), long (L) or short (S) telomeres were crossed in all possible combinations, N/N, N/L, N/S, L/S, S/S, L/L, and telomere length was assessed by Southern blots. Short telomeres were caused by either *yku70Δ* or *mre11Δ* mutations and long telomeres by either *rif1Δ* or *rif2Δ* mutations. To determine how the number of cell divisions affects telomere length, we passaged cultures by streaking for single colonies on agar plates. We estimate a single passage on agar plates corresponds to 25 cell divisions. The cells went through 7 further divisions prior to DNA extraction (see Materials and Methods).

We first examined telomere length using a commonly used Y’ TG probe which detects the large subset of telomeres that contain Y’ repeats (Figure 2A) (Tsubouchi and Ogawa 2000). As expected, this probe showed that Y’ telomeres in wild-type cells were about 1.1 kb, those in *mre11Δ* cells shorter (0.9 kb) and in *rif1*Δ and *rif2*Δ cells longer and more diffuse (1.4 kb) (Figure 2B, lanes 1–6) (Askree et al. 2004) (Wellinger and Zakian 2012). In *mre11Δ/WT* diploids that inherited a set of short telomeres, the Y’-TG probe clearly showed that the telomere lengths had largely normalized by 32 cell divisions (1 passage) (Figure 2B, lane 7), and showed a slight further increase in size during further passage. In contrast, in all diploids containing a *rif1Δ* mutation (*rif1Δ/WT*, *rif1Δ/yku70Δ* and *rif1Δ/rif2Δ*), Y’ telomeres remained longer than normal through 232 divisions (9 passages) (Figure 2B, lanes 10–18). The telomeres of *WT/yku70*Δ diploids, like in *mre11Δ/WT* diploids, rapidly normalized in size (Figure S1C, S1D, lanes 11–13). The telomeres of *rif2Δ/*WT diploids, like those with the *rif1*Δ mutation, remained long for up to 232 divisions (Figure S1A, S1B, lanes 11–16). However, in *rif2*Δ/mre11Δ diploids, Y’ telomeres appeared to approach wild-type size by passage 9 (Figure S1A, S1B, lanes 14–16). This persistence of long telomeres in *rif1*Δ/*WT*, *rif2*Δ/*WT* and *rif1*Δ/*rif2*Δ diploids is presumably due to haploinsufficiency. Interestingly, the *rap1-17* allele affecting the essential protein that binds both Rif1 and Rif2 has semi-dominant effects on telomere length in diploids (Kyrion et al. 1992). Collectively, these data suggest that diploid cells that inherit short telomeres (N/S and S/S) stabilize to wild type telomere lengths comparatively rapidly, whereas long telomeres inherited by N/L, S/L, and L/L diploids either persisted for at least 107 divisions (5 passages).

**Figure 2.**
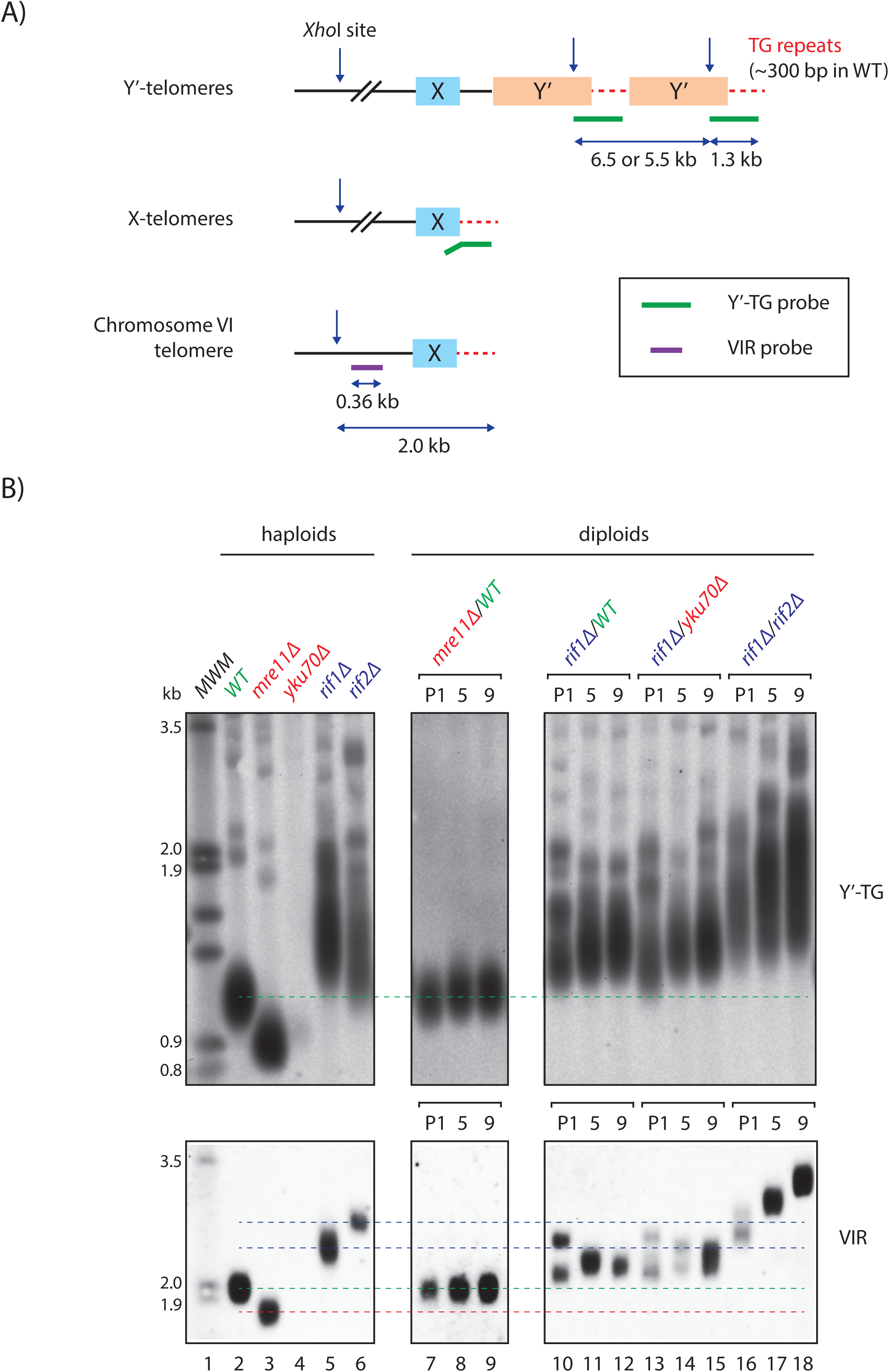
Long telomeres in yeast diploids persist, short telomeres do not. A) Schematic diagram showing *S.cerevisiae* telomeres. The Y’-TG probe anneals to the telomeric end of the Y’-element (∼1 kb) and to terminal TG repeats (∼120 bp). VIR probe anneals upstream of the right X-element of chromosome VI. B) Diploids produced by mating yeast with normal (WT), long (*rif1Δ*, *rif2Δ*) or short (*mre11Δ*, *yku70Δ*) telomeres were examined. More combinations are shown in Figure S1. Telomeres of diploids at passage 1 (32 divisions), 5 (132 divisions) and 9 (232 divisions) were detected by Southern blot using Y’-TG and VIR probes. Sample in lane 4 was poorly digested. Dashed green lines indicate normal (wild-type) telomere length, blue lines indicate long parental haploid telomere length, red lines short parental haploid telomere length.

The Y’ TG probe is useful for telomere blots because it binds to numerous telomeres and gives a strong signal. However, it is not helpful for understanding what happens to individual telomeres in diploid clones. The diffuse signals detected cannot show if specific telomeres are elongated, while others are shortened. To help clarify the picture, a unique VIR probe, which specifically anneals to the right sub-telomeric region of chromosome VI was used (Figure 2A) (Zubko and Lydall 2006). VIR telomere length estimations for Figures 2A and S1 are shown in File S1. The effects of *mre11*Δ (shortening), and *rif1*Δ and *rif2*Δ (elongation) on the length of VIR telomeres in haploids were as expected and mirrored what was seen with the Y’-TG probe (Figure 2B, lanes 2–6). In the *mre11Δ/WT* diploid, by passage 1, VIR telomeres were detected as a single band of WT length (Figure 2B, lane 7). Similar results were observed in *WT/yku70Δ* diploids (Figure S1C, lane 11), confirming the rapid normalization of short telomere lengths in diploids that inherited these from their parents.

The VIR probe was particularly useful to look at telomeres of diploids whose haploid parent(s) had a long telomere mutation. For example, in *rif1*Δ/*WT* diploids at passage 1 two sets of VIR telomeres could clearly be distinguished, the shorter one presumably derived from the wild-type parent and the longer one from the *rif1*Δ parent (Figure 2, lane 10). The long telomere band visible at passage 1, seemed to reduce in length with further passage. We estimate a telomere shortening rate of ∼2 bp/division (from 2.4 to 2.2 kb in a span of 107 divisions, File S1), which is consistent with the rate of degradation of an over-elongated telomere caused by end-replication problem (Marcand et al. 1999) (Figure 2, lane 10 and 11). In the same diploids the shorter VIR telomere was clearly longer than wild-type length at passage 1, and further increased length, presumably due to haploinsufficiency in *rif1Δ/WT* diploids (Figure 2, lanes 10-12). The long and short telomeres ultimately converged to form one tighter band by 232 cell divisions (passage 9), which remained above WT length. Similar patterns of long telomere shortening were seen in *rif1Δ/yku70Δ* diploids (Figure 2, lanes 13–15, File S1). In *mre11Δ/rif2Δ* diploids (Figure S1A, B, lanes 14– 16), the long VIR telomere shortened more noticeably after passage 5 reaching nearly wild-type size by passage 9. These data show that inherited long telomeres can persist for many divisions but may eventually normalize.

Interestingly, in one *rif1Δ/rif2Δ* diploid, VIR telomeres continued to increase in length up to 232 divisions (Figure 2, lanes 16-18). This is consistent with compound haploinsufficiency of *rif1*Δ and *rif2*Δ mutations increasing telomere length. However, the second *rif1Δ/rif2Δ* diploid clone did not illustrate this increase in length (Figure S1B, lanes 23–25), consistent with there being clonal variation. Never the less the telomeres in the second clone were longer than normal, presumably due to haploinsufficiency. More generally, all pairs of diploid clones with identical genotypes, tended to display similar but slightly different patterns of VIR telomere length change with cell divisions (Figure S1).

Taken together our results showed that when diploid yeast cells inherit short telomeres from one parent and normal telomeres from the other parent, telomeres rapidly normalize in length (within 32 mitotic cell divisions). However, in diploid cells that have inherited long telomeres from one parent and normal/short/long telomeres from the other, long telomeres generally persist for more than 32 cell divisions ( at least 100). Long telomeres appear more variable in length and are likely influenced by haploinsufficiency and clonal variation.

### Telomere length evolves slowly in haploids after introduction of long telomere mutations

Since in humans there is evidence that long telomeres can pass down the generations (mitoses and meioses) we wondered if the same were true in yeast. Many of the yeast strains examined in Figure 2 were generated in different laboratories, and there was a lack of clarity about how strains were created and whether, for example, any other mutations were present. Thus, to better understand long telomere inheritance patterns we created new *rif1Δ* and *rif2Δ* mutants by transformation in a clean genetic background.

G418^r^ segments of DNA from *rif1Δ*, *rif2*Δ S288C mutant strains were amplified by PCR and transformed into the W303 WT cells. Two transformant clones of *rif1Δ* and two of *rif2*Δ were selected and passaged up to 35 times (around 900 divisions) (Figure 3). At passage 1, *rif1Δ* mutants had longer telomeres than *rif2Δ* mutants, as previously reported (Wotton and Shore 1997). Also, importantly, our analysis clearly showed that in all four cases VIR telomere length evolved (both increased and decreased) with time. At early passages telomeres grew longer, while at later passages, the occasional reduction in telomere length was observed. The calculated rates of VIR shortening in these instances (lanes 7–8, 15–16, 21–22, 27–28, 29–30) range from 0.7 to 2.2 bp/division (File S1). These rates are consistent with shortening by the end replication problem, and telomerase not being active at extremely long telomeres (Marcand et al. 1999). Alternatively, telomere rapid deletion could have taken place. Y’-telomeres increased in average length in all strains with time, as previously reported (Levy and Blackburn 2004) and as observed in *rap1-18* cells (Kyrion et al. 1992). These results once again demonstrate how single telomere analysis can reveal dynamics that are hidden when examining bulk telomeres.

**Figure 3.**
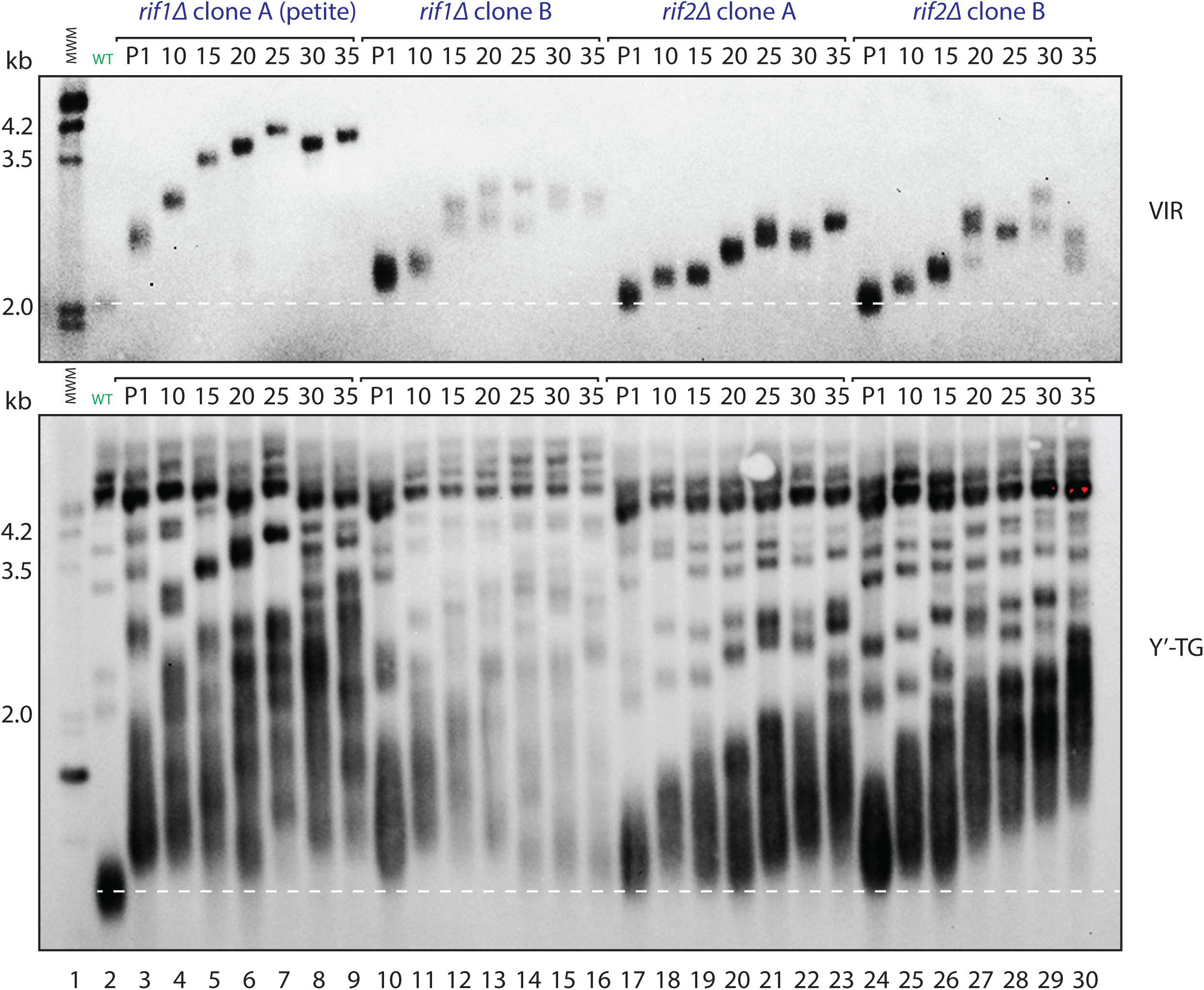
Telomere length of rif1Δ and rif2Δ strains evolves. *RIF1* or *RIF2* genes were disrupted in wild-type cells. Transformants were cloned and telomere length of cells from passages 1 (57 divisions), 10 (281 divisions), 15 (406 divisions), 20 (531 divisions), 25 (656 divisions), 30 (781 divisions) and 35 (906 divisions) were detected as in Figure 2. Dashed white lines indicate normal (wild-type) telomere length. NB at passage 1 *rif1Δ* clone A became petite – a respiratory-deficient mutant with partial or complete loss of mitochondrial DNA (the number of divisions estimated was similar to non-petite clones).

Overall, analysis of VIR telomere length shows that mutations causing long telomeres have effects over many cell divisions, and that the long telomeres eventually generated are unstable, due to telomere rapid deletion or the end replication problem. Moreover, as we had observed in diploids, there is clonal heterogeneity between strains of identical genotypes. Interestingly, we also observe VIR telomere length splitting within one population (Figure 3, lane 6, lanes 12-14 and 27-30), which is unexpected because each passage involves cloning, in principle via a single cell. This may be explained by events like telomere rapid deletion taking place very early on in the growth of a colony from a single cell.

Having obtained the new long telomere mutants with clear provenance, we next used these to follow telomere length changes through the yeast sexual cycle.

### Long telomeres can be inherited independently of the mutations causing long telomeres

To examine the inheritance of long telomeres though meiosis, a newly generated *rif2Δ* strain (Figure 3, clone A, passage 25) was crossed with a WT strain and induced to sporulate. Twelve tetrads were dissected, six produced sets of four viable spores. The spore progeny were scored for mating type, *rif2*Δ status, and telomere length (Figure 4). To simplify our analyses, we classified telomere VIR lengths as being either long or normal (and usually two of each class) even though telomere lengths would be changing in the progeny depending on the *rif2*Δ status and starting length inherited.

**Figure 4.**
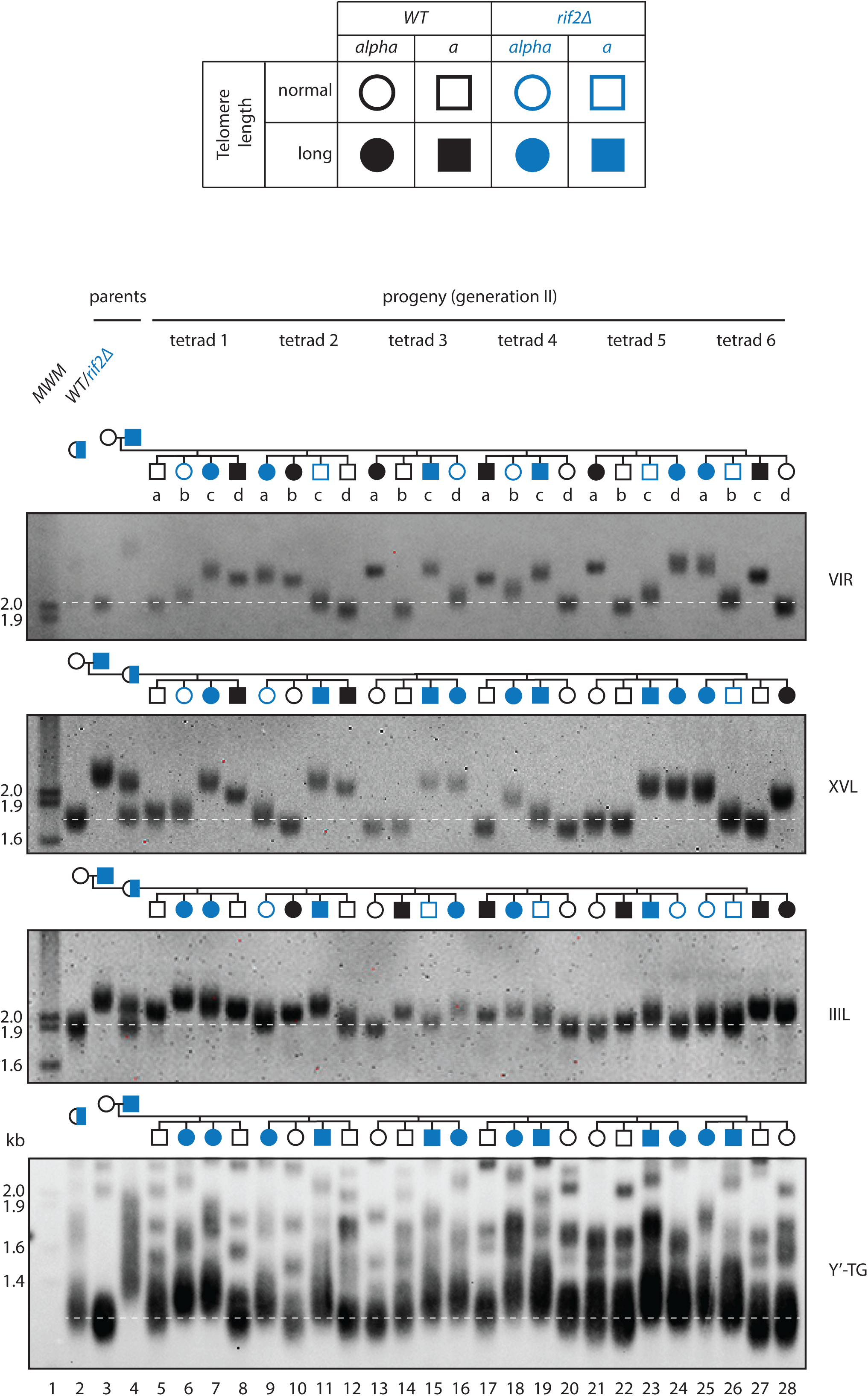
Long telomeres can pass through meiosis independently of mutations causing long telomeres. WT and *rif2Δ* cells (from Figure 3 lane 8, passage 25 (750 divisions)) were mated and sporulated. Tetrads were dissected, and haploids underwent approximately 30 divisions before DNA was prepared. Telomeres were detected as in Figure 2 or using probes to XVL or IIIL. *RIF2* is on chromosome XII and should not be physically linked with any of the telomeres detected. Dashed white lines indicate normal (wild-type) telomere length.

In tetrad 1, we observed a discrepancy between genotype *(rif2*Δ status) and phenotype (VIR telomere length). For example, clone 1b was a *rif2*Δ mutant with normal length telomeres (blue open circle) and clone 1d was wild type with long telomeres (black filled square). A similar pattern was seen in tetrads 2–6, with VIR telomere lengths not correlating with *rif2*Δ status (either blue open symbols, or black filled symbols). In all six tetrads a tetratype segregation pattern was observed between *rif2*Δ and VIR telomere length showing that *rif2*Δ segregates independently of long telomere length.

VIR is reported to be an average length X telomere (Sholes et al. 2022), but to determine if the VIR telomere inheritance patterns we observed were unusual, we examined two other single telomeres. We chose IIIL, because it is longer than average across different strains of *S.cerevisiae* (Sholes et al. 2022; O’Donnell et al. 2023) and XVL as another average sized X telomere. Importantly the *rif2*Δ/*WT* strain transmitted long XVL and IIIL telomeres independently of the *rif2*Δ mutation (either blue open symbols, or black filled symbols)

To systematically assess segregation patterns, we analyzed the six complete tetrads from the cross shown in Figure 4, scoring *rif2*Δ, mating type, and telomere lengths of VIR, XVL, and IIIL. For each pair of markers, we classified the tetrads as parental ditype (PD), non-parental ditype (NPD), or tetratype (TT) (File S2). The predominance of tetratypes when scoring *rif2*Δ status and long telomere length (13 TT, 4 PD, 1 NPD) supports the conclusion that these markers are not genetically linked. Thus, diploid yeast cells carrying a long telomere-inducing mutation can transmit abnormally long telomeres to haploid progeny independently of the mutation that caused telomere elongation, extending earlier qualitative results (Kyrion et al. 1993). Furthermore, different long telomeres segregate independently of one another.

To further validate these findings, an independent cross between a *rif2*Δ strain with longer telomeres (spore 3c from Figure 4) and wild type was performed. Of twelve tetrads dissected, five produced four viable spores (Figure S2). For the *rif2Δ* × VIR long telomere pair, although tetratypes did not predominate, their presence in 2 out of 5 tetrads is inconsistent with there being genetic linkage.

Interestingly analysis of bulk telomeres showed a different pattern. Y’-telomere length inheritance correlated with *RIF2* status: the two *rif2*Δ progeny contain longer telomeres than WT (the absence of blue open symbols or black filled symbols). This shows the effect of the *rif2*Δ mutation on bulk Y’ telomeres masks what may be happening to individual telomeres and/or that there are differences in the effects of Rif2 on X- and Y’-telomeres (Kyrion et al. 1992; Craven and Petes 1999).

In human telomere syndromes, abnormal telomere lengths can be passed down several generations e.g., from grandparent, to parent, to child. To see whether a similar phenomenon occurred in yeast, we examined whether “wild-type” haploid cells, with an abnormally long VIR telomere, could transmit this telomere through another round of meiosis.

A WT cell with long VIR telomeres (Figure 4, clone 3a, lane 13) was mated to a WT cell with normal telomeres and sporulated. It was clear that after this cross some progeny inherited long VIR telomeres (Figure 5A, clones 2b, 2d and 3a lanes 10, 12, 13). Furthermore, when clone 2a (Figure 5A) was further crossed to wild type, long telomeres were once again transmitted to the next generation (Figure 5B).

**Figure 5.**
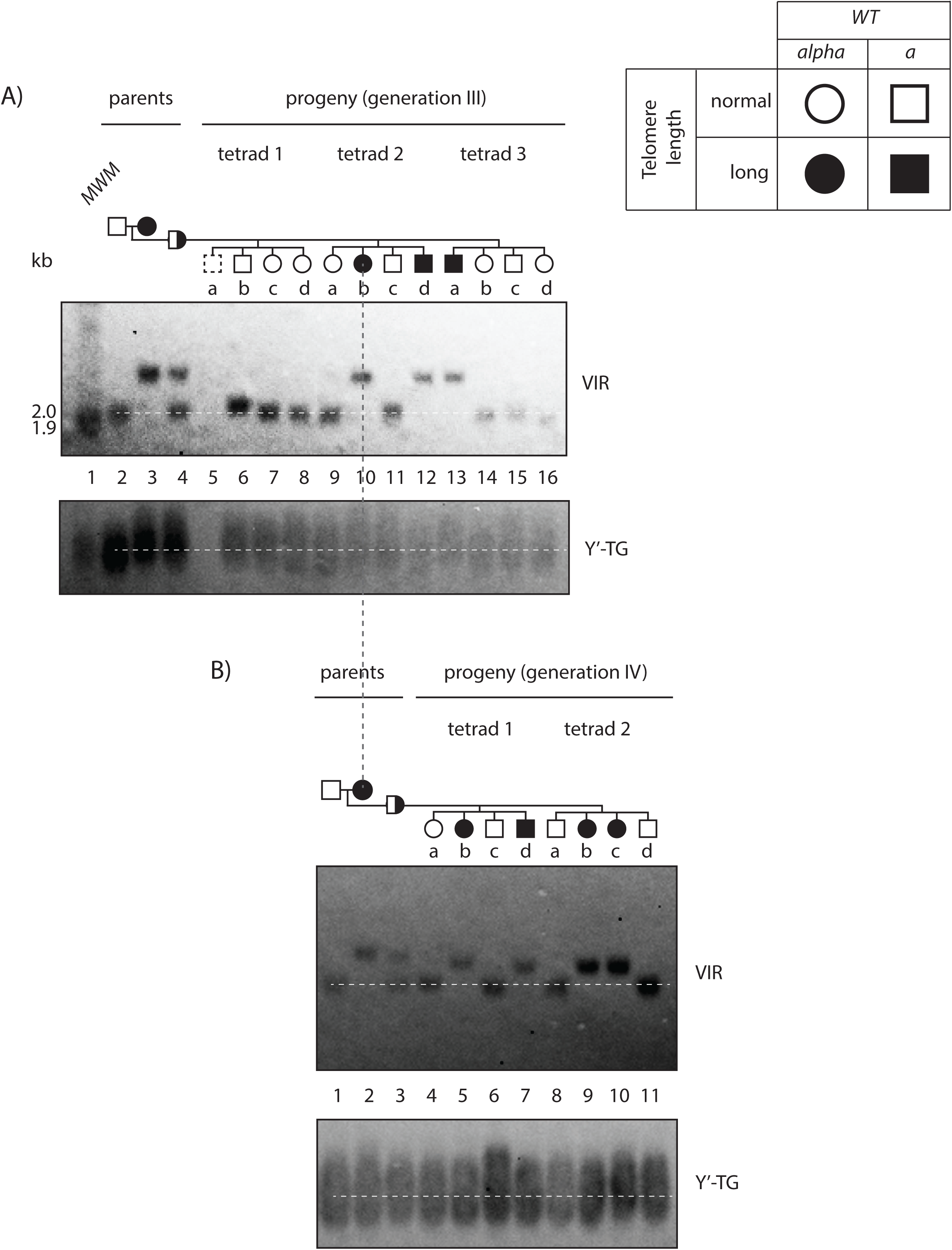
Long telomeres can be inherited through two rounds of meiosis in a WT background. A) WT cells with normal telomeres and WT cells with long telomeres (from Figure 4, clone 3a) were mated and sporulated. Telomeres were detected using Y’-TG and VIR probes. B) WT cells with normal telomeres and WT cells with long telomeres (from Figure 5, clone 2b) were mated and sporulated. Telomeres were detected using Y’-TG and VIR probes. Dashed white lines indicate normal (wild-type) telomere length.

Across the two sexual cycles, VIR telomeres got shorter: from 2.5 kb (clone 3a in Figure 4) to 2.3 kb (clone 2b in Figure 5B), however, the cells never reached the WT VIR length – 2.0 kb. Thus, even after two rounds of meiosis and approximately 110 mitotic cell divisions (see Materials and Methods) in a wild-type genetic background abnormally long VIR telomeres remained. Notably, the rate of VIR telomere shortening was approximately 2 bp/division which aligns with the long VIR shortening rate observed in diploid mitotic cells (Figure 2, Figure S1) and a previously reported rate of shortening of an over-elongated telomere in telomerase-positive cells (Marcand et al. 1999). Interestingly, analysis of tetrad 3 suggested that one elongated telomere had been shortened via recombination event (telomere rapid deletion), since telomeres of three of four progeny were close to normal length while one had very long telomeres. Overall, our experiments clearly show that long telomeres can pass through two yeast sexual cycles/generations in the absence of the mutation that caused the long telomeres. These data also show that persistence of long telomeres in diploids does not depend on any haploinsufficiency.

### Testing the effect of heterozygous short telomere mutations on the shortening of long telomeres in diploids

Long telomeres seem comparatively stable through meiotic and mitotic cell divisions and so we wondered if there might be a way to increase the rate of length normalization. *rif1*Δ and *rif2*Δ mutations appear to show haploinsufficiency effects on telomere length. We hypothesized that mutations that cause short telomeres in haploids might also have haploinsufficient effects, although our data in Figure 2 and Figure S1 show no such effects on short telomeres. Since Mre11, Nmd2, Tel1 and Yku70 are each multifunctional proteins (e.g. with roles in DNA repair, as well as at telomeres) it seemed plausible that some functions may be affected by haploinsufficiency. To test the effects of haploinsufficiency on long telomeres we created diploids that contain long VIR telomeres (from clone 3a in Figure 4) and that were heterozygous *yku70Δ*, *mre11Δ*, *tel1Δ*, or *nmd2Δ* mutations. Four clones (A–D) of each diploid were created and passaged 7 times and telomere length was examined by Southern blotting (Figure S2A–D).

The rates of VIR telomere shortening (bp/division) in different diploids are shown in Figure 6. Consistent with all our earlier experiments there was noticeable clonal heterogeneity in rates of telomere shortening. Although we observed that all four heterozygous mutations seemed to slightly elevate the rate of telomere shortening: *yku70*Δ by 57%, *mre11*Δ by 17%, *tel1*Δ by 24%, and *nmd2*Δ by 40% (File S3), the differences were not statistically significant. Additional replicates would be required to draw statistically robust conclusions regarding the impact of short telomere mutations on long telomere shortening.

**Figure 6.**
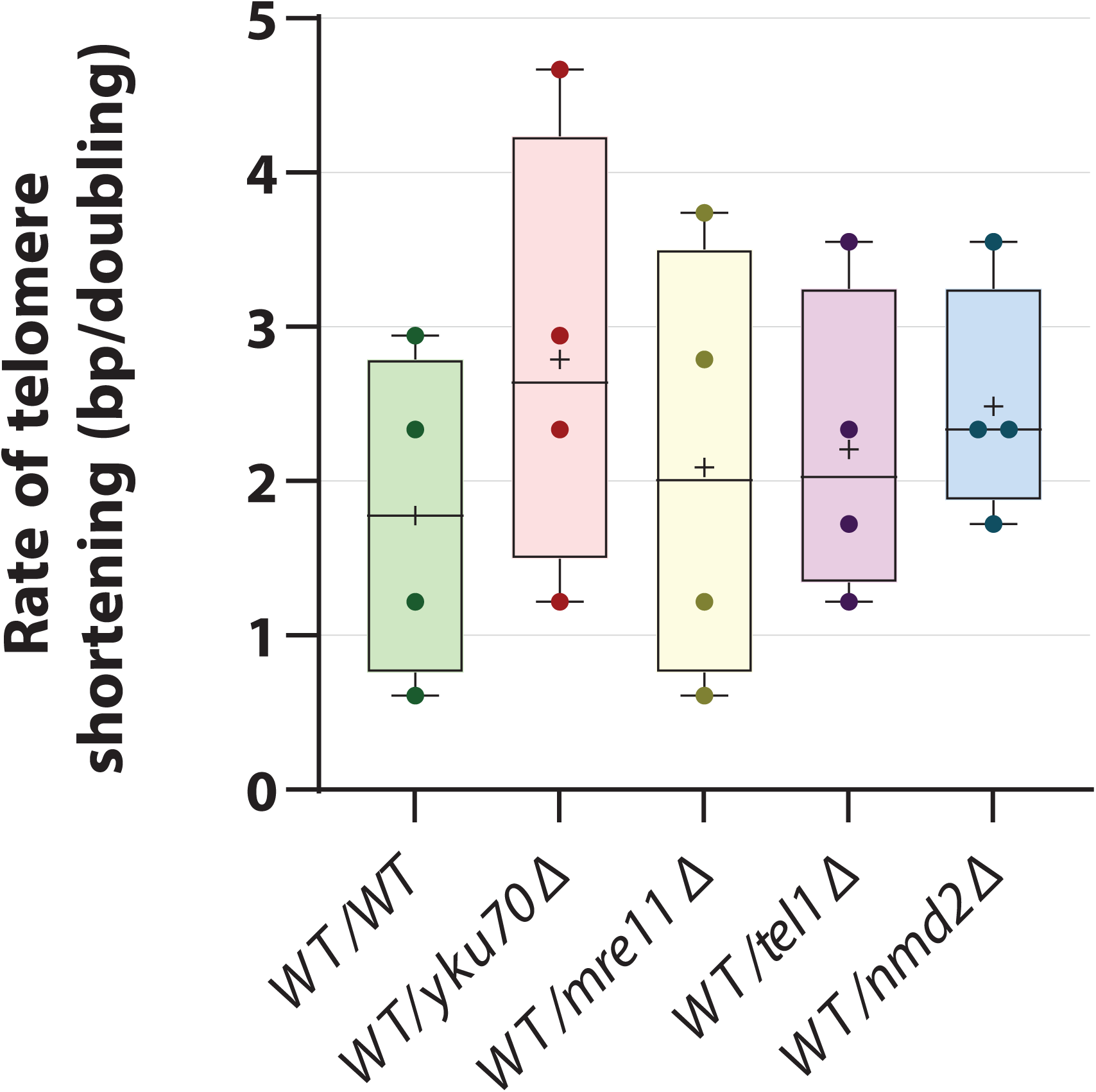
Effect of yku70Δ, mre11Δ, tel1Δ or nmd2Δ mutations on long telomeres. WT cells with long telomeres (clone 3a, Figure 4) were mated with *yku70Δ, mre11Δ, tel1Δ and nmd2Δ* haploids. The rates of telomere shortening were calculated using rate of shortening between passages 0 and 1 (haploid parental cell to zygote derived colony), and passages 1 to 4 of the diploids. Solid round symbols are individual strain rates, plus symbols mean rates and horizonal lines median rates. Telomere length measurements and rate calculations are provided in File S3. Telomere shortening rates were compared across genotypes using one-way ANOVA (p = 0.755).

## Discussion

We have systematically explored telomere length inheritance patterns in the simple single celled eukaryotic model organism budding yeast. The budding yeast model system allows us to switch cells between diploid and haploid states, and to control the number of cell divisions in either state (Figure 1). Another advantage of the yeast experimental model over some other model organisms is that all four products of a single meiosis (spores) are contained in a single ascus and can be isolated and analyzed as a set (a tetrad). For comparison, in female mice, one product of meiosis remains in the egg, while the others are discarded as polar bodies. In male mice, the four meiotic products are viable sperm cells, but any individual sperm cell cannot be connected to its related meiotic products. Our experiments show that even in yeast telomere length inheritance patterns are complex and that this complexity is compounded by significant heterogeneity in pattern between ostensibly genetically identical cells.

Broadly, we draw the following conclusions. Diploids that have inherited short telomeres from one parent rapidly restore telomere length to normal, presumably because telomerase targets short telomeres (Bianchi and Shore 2007). Diploids that have inherited long telomeres from one parent do not rapidly restore telomere length to normal, and long telomeres can persist in diploids and can be passed on through meiosis, to the next generation(s). Furthermore, long telomeres can be transmitted through meioses in the absence of an initial mutation that caused long telomeres. Our analysis also revealed telomere rapid deletion, haploinsufficiency and clonal heterogeneity. It is known that long telomeres in diploids are more susceptible to rapid deletion (recombination) during meiotic cell divisions (Joseph et al. 2005). Consistent with this we saw clear evidence of telomere rapid deletion in one of eleven meiosis we followed (tetrad 3, Figure 5A). The long telomere mutations we examined (*rif1*Δ and *rif2*Δ) showed clear evidence of haploinsufficiency in the diploid state. Finally, ostensibly genetically identical clones, show clear heterogeneity in telomere length inheritance.

Our systematic experiments on long telomere inheritance in yeast are consistent with earlier, less systematic studies. For example, when cells with highly elongated and abnormal telomeres were crossed with wild type cells, all four haploid progeny had highly abnormal telomere lengths (Zubko and Lydall 2006). Similarly, others showed that abnormal long telomeres were transmitted though meiosis (Kyrion et al. 1993; Joseph et al. 2005). Importantly, heritability of abnormal telomere length has also been documented in other well-established model organisms in telomere biology such as mice and zebrafish (reviewed in Henriques and Ferreira 2024). Specifically short telomeres were shown to persist for generations (Hathcock et al. 2002; Hao et al. 2005; Scahill et al. 2017) even in the absence of telomerase mutation that initially caused telomere shortening (Hao et al. 2005). In our experiments in yeast abnormally long telomeres were more stable in diploids than abnormally short telomeres.

There is evidence that human heritable long telomeres are associated with cancer (Horn et al. 2013; Huang et al. 2013; DeBoy et al. 2024). If the human germline behaves similarly to yeast cells, then this suggests that children of a parent carrying long telomeres may inherit long telomeres whether or not they inherit the mutation that causes the long telomeres. Even though these children will also inherit normal length telomeres from a healthy parent, there are several ways in which the inherited long telomeres could affect the risk of cancer. Since telomeric DNA is a challenge for replication fork progression (Sfeir et al. 2009; Olson and Wuttke 2024), longer than normal telomeres would potentially lead to replication fork collapses. This might result in critically short telomeres, triggering genomic instability and carcinogenesis. It is known that telomerase preferentially targets short telomeres. In cells that have inherited a significant number of long telomeres, telomerase will target normal length telomeres to increase all telomere lengths above normal. This dynamic could enhance the individual’s cells proliferative potential and further promote carcinogenesis. Finally, longer telomeres may also affect gene expression from sub-telomeric regions. Overall, it seems plausible that for the management of short and long telomere syndromes, it would be beneficial to assess the length of individual telomeres and across all members of affected families.

## Data availability

All strains used in this study are available upon request. Raw data, including TIFF images of Southern blots, are publicly available at [repository link will be made available].

**Figure S1.** *Set of diploid yeast produced by mating haploids with (WT), long (rif1Δ, rif2Δ) or short (mre11Δ, yku70Δ) telomeres as described in Figure 2.* Telomeres of diploids at passage 1 (32 divisions), 5 (132 divisions) and 9 (232 divisions) were detected by Southern blot using Y’-TG and VIR probes. Red stars indicate poorly digested samples. White lines indicate normal (wild-type) telomere length.

**Figure S2.** *A cross between WT and rif2Δ strains.* WT cells and *rif2Δ* cells with long telomeres (from Figure 4, clone 3c) were mated and sporulated. Telomeres were detected using Y’-TG and VIR probes. Dashed white lines indicate normal (wild-type) telomere length.

**Figure S3.** *Telomere blots showing raw data used in Figure 6.* WT cells with long telomeres (from Figure 4 clone 3a) were mated *yku70Δ, mre11Δ, tel1Δ and nmd2Δ* haploids. Four diploid clones (A–D) from each mating were passaged and examined by Southern blot. Telomeres at passage 1 (32 divisions), 4 (107 divisions) and 7 (182 divisions) were detected using Y’-TG and VIR probes. Dashed white lines indicate normal (wild-type) telomere length. A single haploid wild type clone, with a long VIR telomere was used as a control on all four blots (lanes 27–29).

**File S1.** *Estimations of telomere lengths for Figures 2–5, S1–S2*

**File S2.** *Analysis of segregation patterns of rif2Δ, mating type and telomere lengths of VIR, XVL, and IIIL for Figures 4 and S2.*

**File S3.** *Estimations of VIR telomere lengths and VIR shortening rates for Figure S3.*

## Acknowledgements

We would like to thank Sveta Makovets for input and her comments on the manuscript. We thank Peter Banks, Alessandro Bianchi, Martin Kupiec, Ed Louis, Art Lustig, Laura Maringele, Simon Whitehall for advice and/or yeast strains. We thank Mundy Wellinger and Gabriela Teplitz for advice on Southern blot probes.

